# The aryl hydrocarbon receptor regulates epidermal differentiation through transient activation of TFAP2A

**DOI:** 10.1101/2023.06.07.544032

**Authors:** Jos P.H. Smits, Jieqiong Qu, Felicitas Pardow, Noa J.M. van den Brink, Diana Rodijk-Olthuis, Ivonne M.J.J. van Vlijmen-Willems, Simon J. van Heeringen, Patrick L.J.M. Zeeuwen, Joost Schalkwijk, Huiqing Zhou, Ellen H. van den Bogaard

## Abstract

The aryl hydrocarbon receptor (AHR) is an evolutionary conserved environmental sensor identified as indispensable regulator of epithelial homeostasis and barrier organ function. Molecular signaling cascade and target genes upon AHR activation and their contribution to cell and tissue function are however not fully understood. Multi-omics analyses using human skin keratinocytes revealed that, upon ligand activation, AHR binds open chromatin to induce expression of transcription factors (TFs), e.g., Transcription Factor AP-2α (TFAP2A), as a swift response to environmental stimuli. The terminal differentiation program including upregulation of barrier genes, filaggrin and keratins, was mediated by TFAP2A as a secondary response to AHR activation. The role of AHR-TFAP2A axis in controlling keratinocyte terminal differentiation for proper barrier formation was further confirmed using CRISPR/Cas9 in human epidermal equivalents. Overall, the study provides novel insights into the molecular mechanism behind AHR-mediated barrier function and potential novel targets for the treatment of skin barrier diseases.

## Introduction

The skin, being an important barrier organ, plays a major role in protecting and fostering the life it encloses. Within the ever-renewing epidermis, keratinocytes are the predominant cell type, accounting for 95% of epidermal cells^1^. The continuous renewing of the epidermis is highly dependent on the delicate balance between keratinocyte proliferation and differentiation. During epidermal development, basal stem cells give rise to daughter cells which undergo a coordinated program of cell cycle arrest, upward migration, and terminal differentiation. Maintaining the integrity of the epidermis is essential for skin homeostasis and protection of the host against infections, allergens, UV radiation and other external threats through host defense, and physical, chemical, and immunological barrier mechanisms^2^. As such, a compromised epidermal barrier is a prominent feature of common inflammatory skin diseases, like atopic dermatitis and psoriasis^3,4^. In healthy skin, epidermal homeostasis is tightly controlled through a set of essential transcription factors (TFs), *e.g.*, TP63, AP1, and the aryl hydrocarbon receptor (AHR)^5–7^.

AHR is a TF that is considered a sensor of environmental, microbial, metabolic, and endogenous cues. Depending on the specific activating ligand, AHR activation can cascade into a response ranging from highly toxic to therapeutic^8–10^. AHR is involved in many biological processes, from cellular proliferation and differentiation to immune responses both innate and adaptive of origin. Upon activation, AHR translocates from the cytoplasm to the nucleus, where it dimerizes with AHR nuclear transporter (ARNT) to bind to its cognate DNA consensus sequence (5′-TNGCGTG-3′) known as the xenobiotic response element (XRE) and regulates gene transcription^8,11^. Certain AHR-activating ligands are highly toxic, e.g., high-affinity environmental pollutant dioxins (e.g., 2,3,7,8-Tetrachlorodibenzo-*p*-dioxin (TCDD)). TCDD has an extremely long half-life resulting in prolonged and uncontrolled AHR activation^12^, while other AHR ligands are rapidly degraded and considered of more physiological importance, e.g., 6-formylindolo[3,2-b]carbazole (FICZ), which is generated upon UV radiation of keratinocytes^13^.

Over the years, we have gained better understanding of the effects of AHR activation on inflammatory skin conditions since the discovery of AHR activation as the working mechanism of coal tar (CT) ointment that was used for psoriasis and atopic dermatitis treatment^14–16^. These insights sparked the global interest in therapeutics that target the AHR in skin diseases and beyond, and led to the registration of Tapinarof, an AHR ligand, for psoriasis^17,18^. Phase 3 clinical trials in atopic dermatitis are ongoing (NCT05032859). Other AHR ligands with similar biological implications, including carboxamide and indazole derivatives, have also been studied for their therapeutic anti-inflammatory and barrier promoting potential ^19–23^.

At the molecular level, mainly four groups of genes are known to be targeted by AHR in the skin. Firstly, a battery of xenobiotic metabolizing enzymes (XMEs), including cytochrome P450 monooxygenases (P450s), e.g., *CYP1A1*^24^; secondly, genes involved in keratinocytes differentiation^25^, e.g., *filaggrin* and *involucrin*^14,26-28^; thirdly, genes related to host defense, e.g., the antimicrobial peptide (AMP) families of *S100* genes, *late cornified envelope* (*LCE*) genes, and *peptidase inhibitor* (*PI*)*3*, amongst others^16,29^; and finally, genes related to immunity, e.g., the inflammatory cytokines *Interleukin (IL)-1β*, *IL-6, CXCL5, CCL20* and *IL-10*^30–32^. Hence, AHR activation is found to increase epidermal differentiation and barrier formation^29,33-35^, and dampen skin inflammation^36^. However, the sequence and dynamics of the molecular events and other players involved through which AHR mediates these effects are poorly understood. In this study, we aim to characterize regulatory cascade upon AHR activation in human keratinocytes. Through transcriptomic and epigenomic analyses, we identified a hitherto unrecognized AHR-TFAP2A axis that regulates epidermal keratinocyte terminal differentiation and skin barrier formation.

## Results

### AHR activation results in distinct early and late transcriptional programs

To characterize the gene expression pattern upon AHR activation in keratinocytes, we performed RNA-sequencing on keratinocytes either treated with TCDD or coal tar (CT), two AHR model ligands, for short term (2 h) and longer exposure duration (24 h). Principle component analysis (PCA) showed transcriptome alterations in ligand treated samples already after 2 h of treatment, indicating that ligand exposure results in swift AHR activation and transcription regulation (Fig. 1a). The differences became increasingly apparent between 2 h and 24 h of ligand treatment, indicated as the major change through PC1 axis (71% variance). Differences between TCDD and CT treated samples were minor as they closely clustered in the PCA plot, indicating that regulatory events downstream of AHR activation are similar in both treatment conditions. *CYP1A1* and *CYP1B1*, target genes in canonical AHR signaling, showed consistent up-regulation upon both ligand treatments, more significantly after 24h treatment (Fig. 1b). Their gene expression was validated with qPCR (Fig. 1c). These observations indicate that TCDD and CT treatment activate AHR signaling pathways through a similar pool of genes within 24 h, and we therefore focused on the common mechanism shared between TCDD and CT treatment in subsequent analyses, hereafter referred to as “ligand-treatment”.

**Fig. 1.**
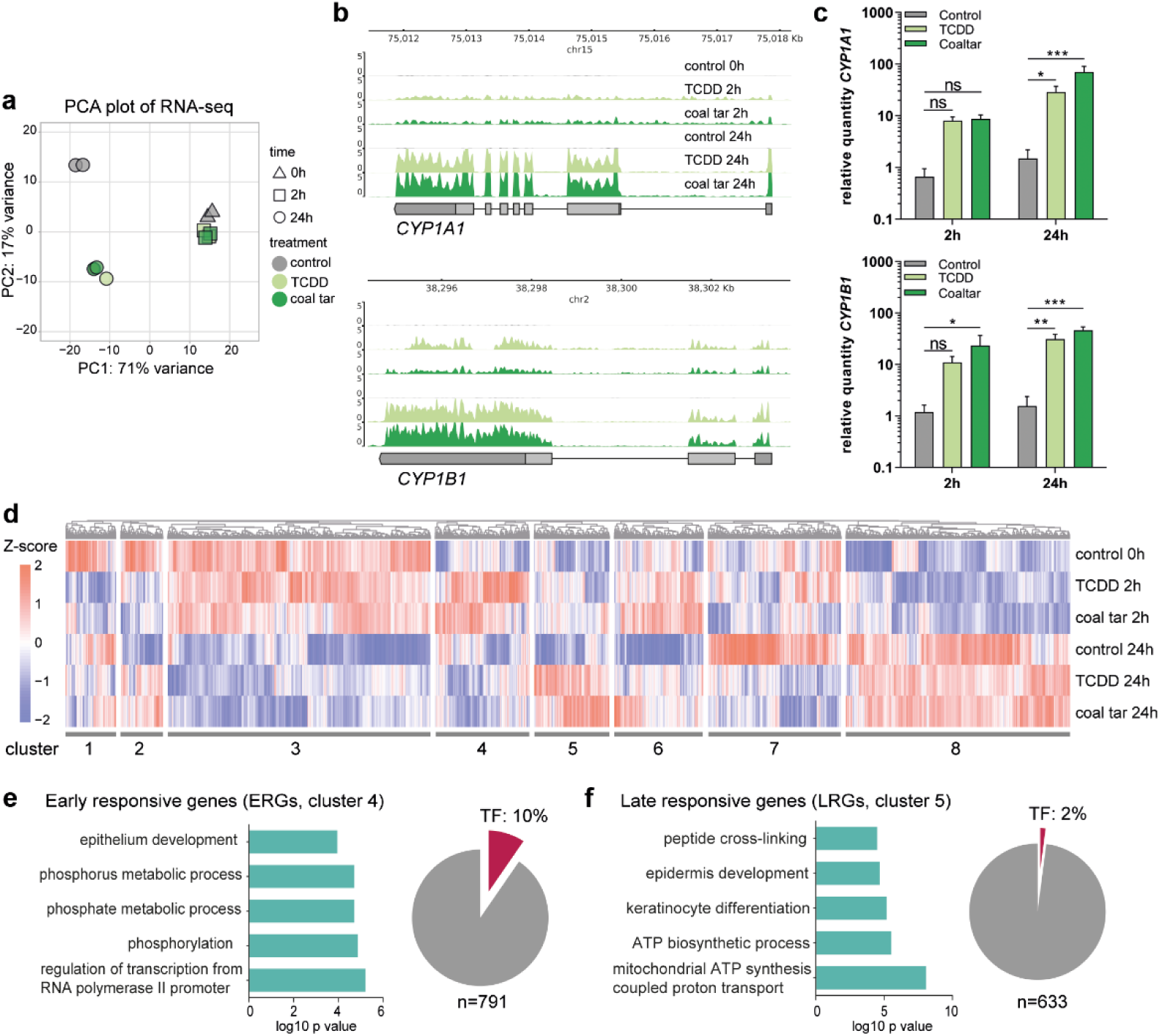
AHR activation results in distinct early and late response. a. Principal component analysis (PCA) of RNA-seq data. **b** Genome browser screenshots of *CYP1A1* and *CYP1B1* on RNA-seq tracks. **c** RT-qPCR validation of *CYP1A1* and *CYP1B1*. Data shown as mean ±SEM, N=5 technical replicates, two-way ANOVA. **d** Hierarchical clustering of differentially expressed genes (p value <0.05). Z-score was calculated based on log10 (FPKM+0.01) of each gene. **e** GO annotation of ERGs, accompanied by a pie chart showing the number and percentage of TFs within the **f** GO annotation of LRGs, accompanied by a pie chart showing the number and percentage of TFs within the cluster.

Next, we identified differentially expressed genes (DEGs, adjusted p value <0.05) between the control and at both 2 h and 24 h of ligand treatment. In total, 8160 DEGs were grouped into eight hierarchical clusters according to the gene expression dynamics at different time points after ligand treatment (Fig. 1d, Table 1, and Supplementary Data 1). Clusters 1 and 2 show early downregulation upon ligand-treatment, with no apparent late effects or dampened downregulation after 24 h of treatment, respectively. Cluster 3 and cluster 8 comprised the majority of DEGs but their gene expression was unrelated to AHR ligand treatment and mainly affected by the keratinocyte differentiation itself. Genes from cluster 3 are mainly associated with gene ontology (GO) term ‘cell cycle’ and genes from cluster 8 are involved in ‘translation’. Importantly, genes in cluster 4 showed up-regulated expression after 2 h ligand treatment and are involved in the processes of ‘phosphorylation’ and ‘epithelium development’, e.g., *NOTCH2*, *JUN*, *TFAP2A*, *KRT4*, and *POU3F1*. In contrast, genes in cluster 5 showed late up-regulated expression only after 24 h of ligand treatment and mainly contribute to ‘keratinocyte differentiation’, e.g., *FLG* and *IVL*, and ‘oxidation-reduction process’, e.g., *HYAL1* and *CYCS*. Clusters 6 contains genes that are slightly upregulated early after ligand treatment. These genes appear downregulated at 24 h in control, probably due to differentiation, while ligand treatment at this timepoint dampens the downregulation. Cluster 7 contains genes that are downregulated 24 h after treatment initiation. Interestingly, there was no distinct cluster of genes that showed continuous up-regulation or down-regulation at 2 h and 24 h after TCDD and CT treatment. This highlights the dynamics of AHR signaling in primary cells, rather than the reported continuous signaling in (cancer) cell lines^37^.

**Table 1:** Hierarchical clusters with significantly DE genes of ligand-treated keratinocytes

| Cluster | Cluster size | Gene expression dynamics upon AHR ligand treatment | Most significant GO term | Example genes |
| --- | --- | --- | --- | --- |
| 1 | 425 genes | Early (downregulated) responsive, with no apparent late effects | GO:0070647 protein modification by small protein conjugation or removal | *BBS2, KIF3A* |
| 2 | 355 genes | Early (downregulated) responsive. Genes appear downregulated in 24h control while ligand treatment dampens this downregulation | GO:0051301 cell division | *CDK1* |
| 3 | 2207 genes | Non-responsive to treatment | GO:0007049 cell cycle | *E2F1* |
| 4 | 791 genes | Early (upregulated) responsive (ERGs) | GO:0060429 epithelium development | *NOTCH2, JUN, TFAP2A, KRT4, POU3F1* |
| 5 | 633 genes | Late (upregulated) responsive (LRGs) | GO:0008544 epidermis development | *FLG, IVL* |
| 6 | 748 genes | Early (slight upregulated) responsive. Genes appear downregulated in 24h control while ligand treatment dampens this downregulation | GO:0007049 cell cycle | *HIST1H4L, POLR2F* |
| 7 | 1118 genes | Late (downregulated) responsive genes | GO:0051252 regulation of RNA metabolic process | *HKR1, FOXO3, TGFB2* |
| 8 | 1883 genes | Non-responsive to treatment | GO:0002181 cytoplasmic translation | *RPL18* |

To dissect the molecular events upon AHR activation, we continued to focus on clusters of ‘early-responsive genes (ERGs)’ (cluster 4, Fig. 1e, upregulation 2h after ligand treatment) and ‘late-responsive genes (LRGs)’ (cluster 5, Fig. 1f, upregulation 24h after ligand treatment). The separation of ERGs and LRGs suggests a different regulatory mechanism of AHR signaling between early and late responses. This observation led us to hypothesize that the early and late responses are potentially linked via TFs in ERGs that activate transcription of LRGs. Indeed, among the 8160 DEGs, 558 genes were classified as TFs and 76 TFs out of 791 genes (10%, hypergeometric p value = 0.001) were found in ERGs, e.g., *HES1*, *HES2*, *FOSL1, JUN*, *TFAP2A*, and *SOX4*. In contrast, LRGs did not show a significant enrichment of TFs (13 TFs in 633 genes, e.g., *GRHL1* and *STAT6*, hypergeometric p-value = 1.2)(Fig. 1e, f, Supplementary Data 1). This clear enrichment of TFs in ERGs in contrast to LRGs supports our hypothesis that the up-regulated TFs among ERGs regulated the expression of LRGs, including the expression of epidermal differentiation genes.

### AHR activation promotes dynamic alterations of the enhancer landscape

To identify AHR target genes, including TFs, we set out to first map enhancers bound by AHR. Being a receptor of environmental cues, AHR was expected to bind to chromatin in a swift and transient manner, and therefore we first performed AHR targeted chromatin immunoprecipitation followed by qPCR (ChIP-qPCR) to determine the binding time frame. At 30 min of ligand treatment, AHR binding signals were detected at the loci of the known AHR target gene *CYP1A2* (Fig. 2a). Such fast binding of AHR was consistent with the translocation of AHR from the cytoplasm to the nucleus as shown by immunofluorescent staining after 30 min of ligand treatment (Fig. 2b). Notably, the AHR binding signals decreased after 90 min of ligand treatment (Fig. 2a), confirming the transient character of AHR interaction to its target loci. The dynamic targeting of the genome by AHR in primary keratinocytes is consistent with our observations on gene expression changes upon AHR activation (Fig. 1d).

**Fig. 2.**
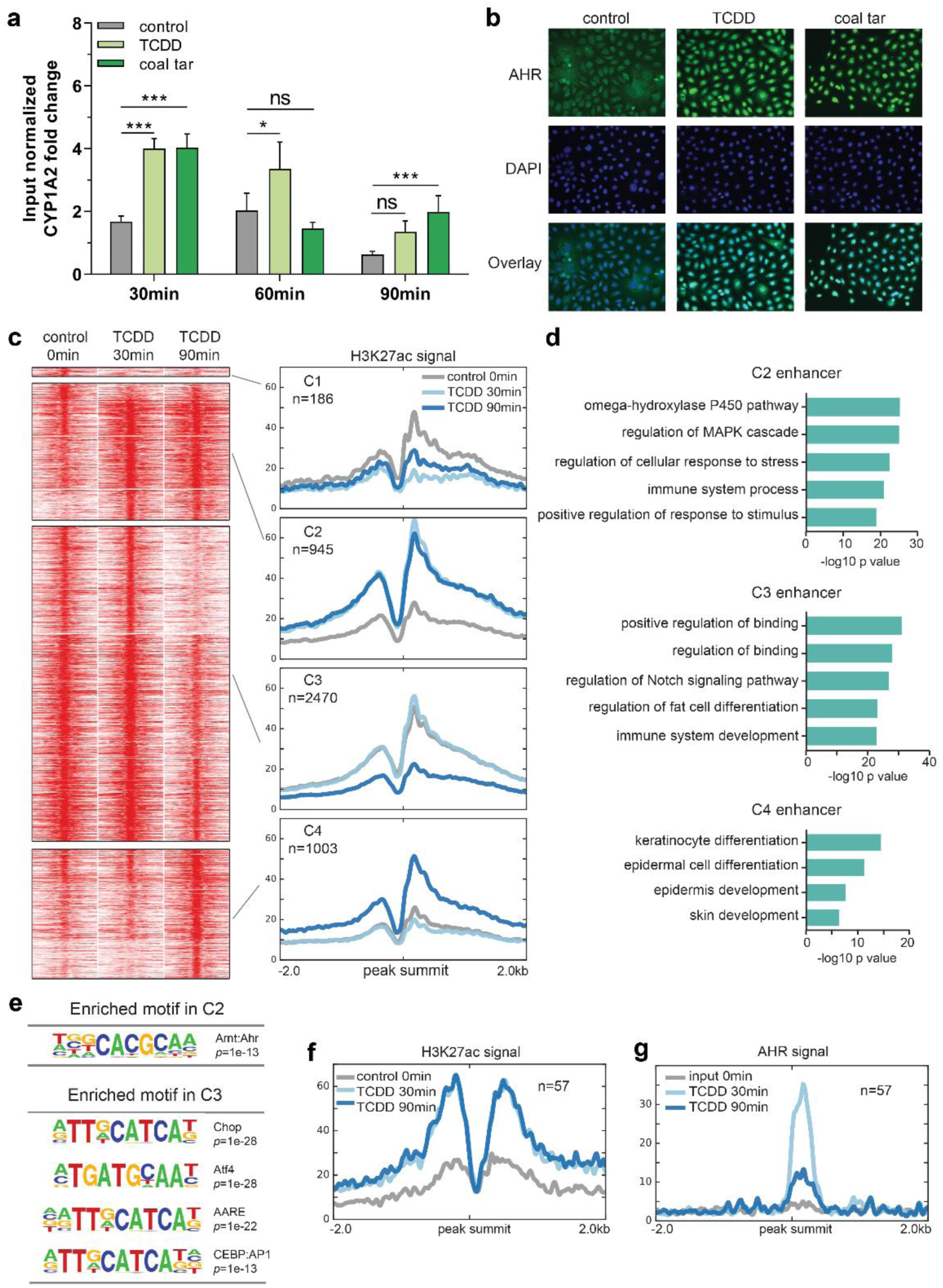
AHR activation leads to enhancer dynamics. a. AHR ChIP-RT-qPCR performed at the loci of *CYP1A2* (as a positive control) at different time point after ligand treatment. Input normalized fold change is relative to both input DNA and negative control loci (chr11). Data shown as mean ±SEM, N=6 technical replicates, two-way ANOVA. **b** AHR translocation from the cytoplasm to the nucleus after 30min of ligand treatment. **c** Clustering of the dynamic enhancers upon AHR activation. Heat maps and band plots are shown in a 4-kb window with summits of enhancers in the middle. Color intensity in heat maps represents normalized read counts. In the band plots, the median enrichment was shown. **d** GO annotation of enhancers in C2, C3 and C4. **e** Significantly enriched motifs found in C2 and C3 of dynamic enhancers shown in **c**. **f** Band plot showing the quantification H3K27ac ChIP-seq signals at AHR binding sites upon ligand treatment. **g** Band plot showing the quantification AHR ChIP-seq signals at AHR binding sites upon ligand treatment.

Since we consistently observed similar gene expression and AHR binding following both TCDD and CT treatments, we continued our experiments with only TCDD stimulation to model AHR activation. To identify AHR-responsive enhancers that are involved in gene activation, we performed H3K27ac ChIP-sequencing after TCDD treatment for 30 and 90 min. Clustering of enhancer regions based on H3K27ac signals gave rise to four clusters consisting of 4,604 enhancers (Fig. 2c and Supplementary Data 2). Subsequently, motif analysis was performed to predict TFs that potentially bind to these enhancers (Fig. 2e, Supplementary Data 2). Among the four clusters, only cluster 1 (shown as C1) containing a small number (186) of enhancer regions showed decreased activity upon AHR activation by TCDD at 30 min (Fig. 2c), and motif analysis did not yield statistically enriched TF motifs (Supplementary Data 2). Cluster 2 (C2) represents 945 enhancer regions that showed a reasonable level of H3K27ac signals in the control (0 min) and increased signals at 30 min of TCDD treatment. The H3K27ac signals remained high after 90 min. Genes nearby these enhancers are mainly involved in ‘omega-hydroxylase P450 pathway’ shown by the GO analysis (Fig. 2d), and contain many known AHR targets, such as *CYP1A1* and *CYP1A2*. TF motif analysis showed that the AHR motif was the only highly enriched motif in C2, indicating that this cluster of enhancers are likely directly bound by AHR (Fig. 2e, Supplementary Data 2). Cluster 3 (C3) contains 2470 enhancer regions maintaining high signals at 0 and 30 min, which decreased after 90 min of TCDD treatment. Genes nearby these enhancers are mainly involved in ‘regulation of Notch signaling pathway’, e.g., *BMP7*, *HES1*, *JAG1*, and ‘immune system development’, e.g., *BCL6* and *CD28* (Fig. 2d). CHOP, ATF4, AARE and CEBP binding motifs from the AP-1 motif family were enriched in C3 enhancers (Fig. 2e, Supplementary Data 2). The last cluster, cluster 4 (C4), consisted of 1003 enhancers showing higher activity only after 90 min of TCDD treatment, with nearby genes being predominantly involved in ‘keratinocyte differentiation’, e.g., *FLG* and *HRNR*. This cluster did not contain significantly enriched TF binding motifs (Supplementary Data 2).

To confirm the motif analysis of C2 in which the AHR motif was enriched, we performed AHR ChIP-sequencing with TCDD treatment and obtained 57 AHR binding sites (adjusted p value = 1e-4, Supplementary Data 3). When examining H3K27ac signals at AHR binding sites, we observed persistent H3K27ac signals at both 30 min and 90 min of treatments (Fig. 2f), fully consistent with C2 cluster enhancer signals (Fig. 2c), confirming this cluster of enhancers being direct targets of AHR. Of note, the apparent H3K27ac signals at most of the C2 cluster enhancers in the control without ligands indicate that AHR binds to open chromatin regions. At the same time, AHR binding signals peaked at 30 min and went down after 90 min of treatment (Fig. 2g), in line with the transient AHR binding observed from ChIP-qPCR analysis of the CYP1A2 locus (Fig. 2a).

In summary, these data demonstrate a transient nature of AHR-enhancer binding, leading to early activation of enhancer targets (C2) and the late activation of enhancers near epidermal differentiation genes (C4). This distinct activation scheme is consistent with the temporal divided expression pattern of ERGs and LRGs (Fig. 1).

### Transcription factor AP-2 Alpha (TFAP2A) is direct target of AHR

To confirm our hypothesis that AHR-controlled TFs among ERGs regulate keratinocyte differentiation program as the secondary response to AHR activation, and to identify such candidate TFs, we integrated the RNA-sequencing, AHR ChIP-sequencing and H3K27ac ChIP-sequencing data. We set the criteria of such intermediate TFs to: exhibiting up-regulated gene expression upon AHR activation by ligands (cluster 4 in Fig. 1d) and denoted by a nearby AHR-bound active enhancer as indicated by AHR and H3k27ac ChIP-sequencing signals. We identified Transcription Factor AP-2 Alpha (TFAP2A), known to play a role in keratinocyte differentiation^38,39^, to fit this profile. TFAP2A is among ERGs and has an intronic AHR-bound enhancer with a high H3K27 signal (Fig. 3a). AHR binding at this locus was validated by ChIP-qPCR (Fig. 3b), establishing TFAP2A as a likely direct AHR target.

**Fig. 3.**
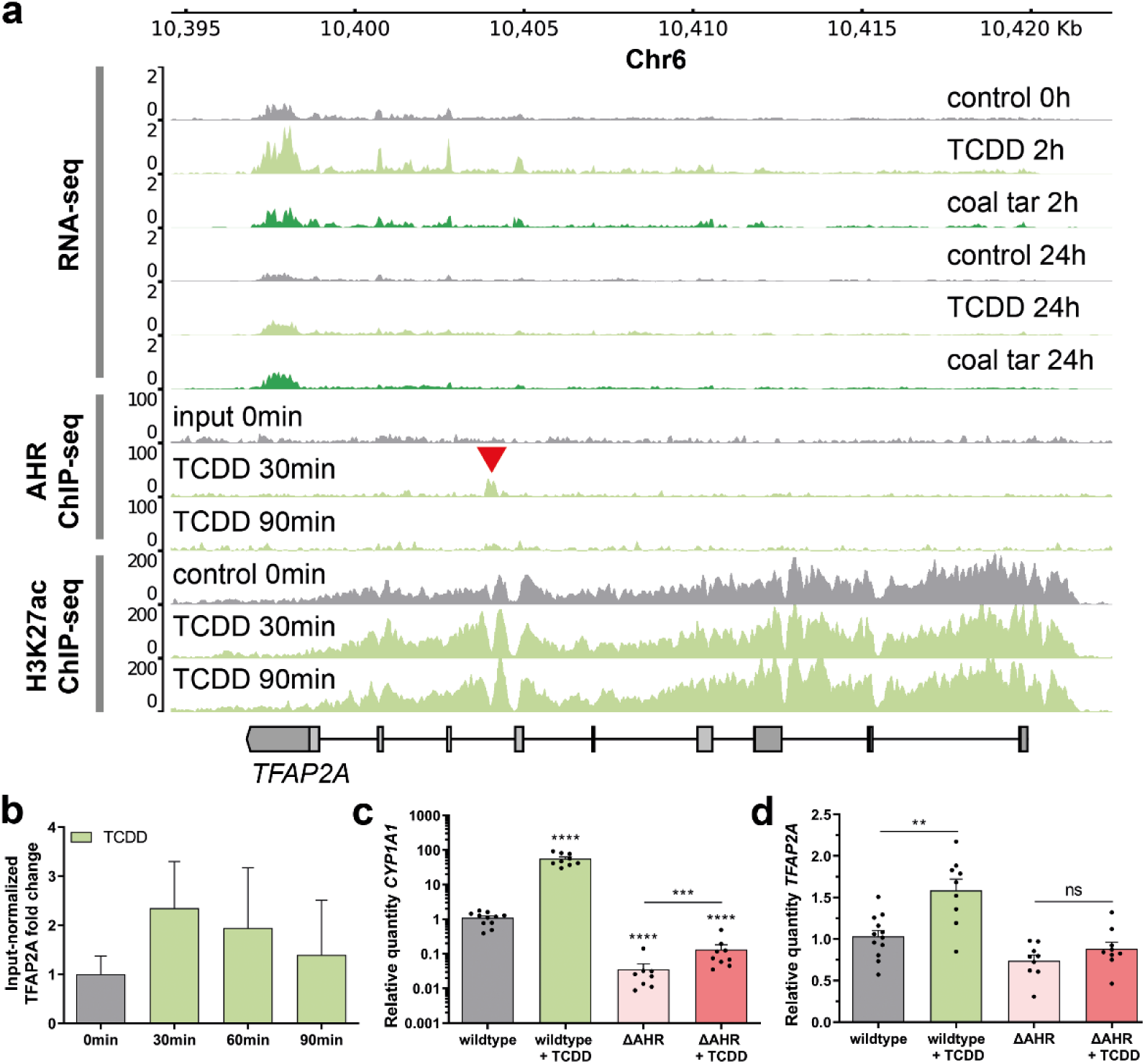
AHR targets TFAP2A in the early response to AHR ligands. a. Genome browser screenshots of the *TFAP2A* coding region show RNA-seq, AHR ChIP-seq, and H3K27ac ChIP-seq tracks upon treatment with coal tar and TCDD. Red arrow indicates AHR binding site within the *TFAP2A* locus. **b** AHR ChIP-RT-qPCR validation at the loci of *TFAP2A* at different time point after ligand treatment. Input normalized fold change is relative to both input DNA and negative control loci (chr11). Data are shown as mean ±SEM, N=2 technical replicates. **c, d** Knockout of AHR (ΔAHR) is accompanied by the loss of *CYP1A1* (as classical AHR target) and *TFAP2A* expression. Data shown as mean ±SEM, N> 5 technical replicates, one-way ANOVA.

To functionally validate whether TFAP2A is a primary AHR target, clonal homozygous AHR knockout (ΔAHR) keratinocytes were generated using CRISPR/Cas9 in the immortalized N/TERT-2G keratinocyte cell line^40^. After clonal expansion of the knockout pool, a full ΔAHR clonal keratinocyte cell line was identified using PCR and subsequent Sanger sequencing. On both alleles one nucleotide was deleted resulting in a frameshift after 76 amino acids, and an early stop codon that translates to a loss-of-function truncated AHR protein. As expected, the expression of a known AHR target gene *CYP1A1* was significantly lower in ΔAHR keratinocytes than that in wildtype cells, and *CYP1A1* expression was not enhanced upon TCDD treatment in ΔAHR keratinocytes, as it was the case in wildtype cells (Fig. 3c). Importantly, ΔAHR keratinocytes showed a loss of target gene expression and TCDD treatment of ΔAHR keratinocytes did not increase the *TFAP2A* expression level, in contrast to the enhanced expression of *AHR* wildtype keratinocytes (Fig. 3d), which is in line with our notion that *TFAP2A* is indeed an AHR direct target gene.

### AHR-TFAP2A axis controls the epidermal differentiation program

Next, we investigated the contribution of TFAP2A activation in AHR-mediated keratinocyte differentiation. We knocked down *TFAP2A* in monolayer primary keratinocyte cultures using siRNAs (52% knockdown compared to siControl; Fig. 4a, b), treated *TFAP2A* knockdown keratinocytes with TCDD for 24h to activate AHR signaling, performed RNA-sequencing analysis, and detected 435 genes that were differentially expressed between TCDD-treated siCtrl and siTFAP2A (Supplementary Data 4). To identify TFAP2A-mediated AHR signaling, we examined the effect of *TFAP2A* knockdown on TCDD-induced gene expression and compared them to the earlier identified panel of AHR-responsive genes (from Fig. 1d, all clusters, 3084 DEGs in total, 1976 genes up- and 1108 genes downregulated). Among the 435 differentially expressed genes upon TFAP2A knockout, 214 are overlapping with the identified 3084 AHR-responsive genes. The overlapping genes were clustered according to their expression patterns (Fig. 4c, Supplementary Data 4). Clusters 1 and 2 (18 and 40 genes, respectively) contain genes that are downregulated by TCDD and remain downregulated (cluster 1) or become upregulated (cluster 2) upon TCDD treatment in *TFAP2A* knockdown condition. Cluster 3 and 4 (57 and 99 genes, respectively) contain genes that are upregulated by TCDD treatment and remain upregulated in both conditions (cluster 3) or become downregulated upon TCDD treatment in *TFAP2A* knockdown condition (cluster 4). Because cluster 4 genes were induced by TCDD and the induction was abolished by *TFAP2A* knockdown, these genes were marked as a TFAP2A-mediated AHR response genes. Cluster 4 was found to be enriched for LRGs (mean fold enrichment 20.8, hypergeometric p-value 7.11e-47), including *IVL*, several *S100* genes, and *SPRR* genes that are involved in terminal differentiation of epidermal keratinocytes. In line with this, functional annotation of the genes in cluster 4 resulted in 57 significantly enriched GO terms, such as ‘epidermis development’ and ‘keratinocyte differentiation’ (Fig. 4d).

**Fig. 4.**
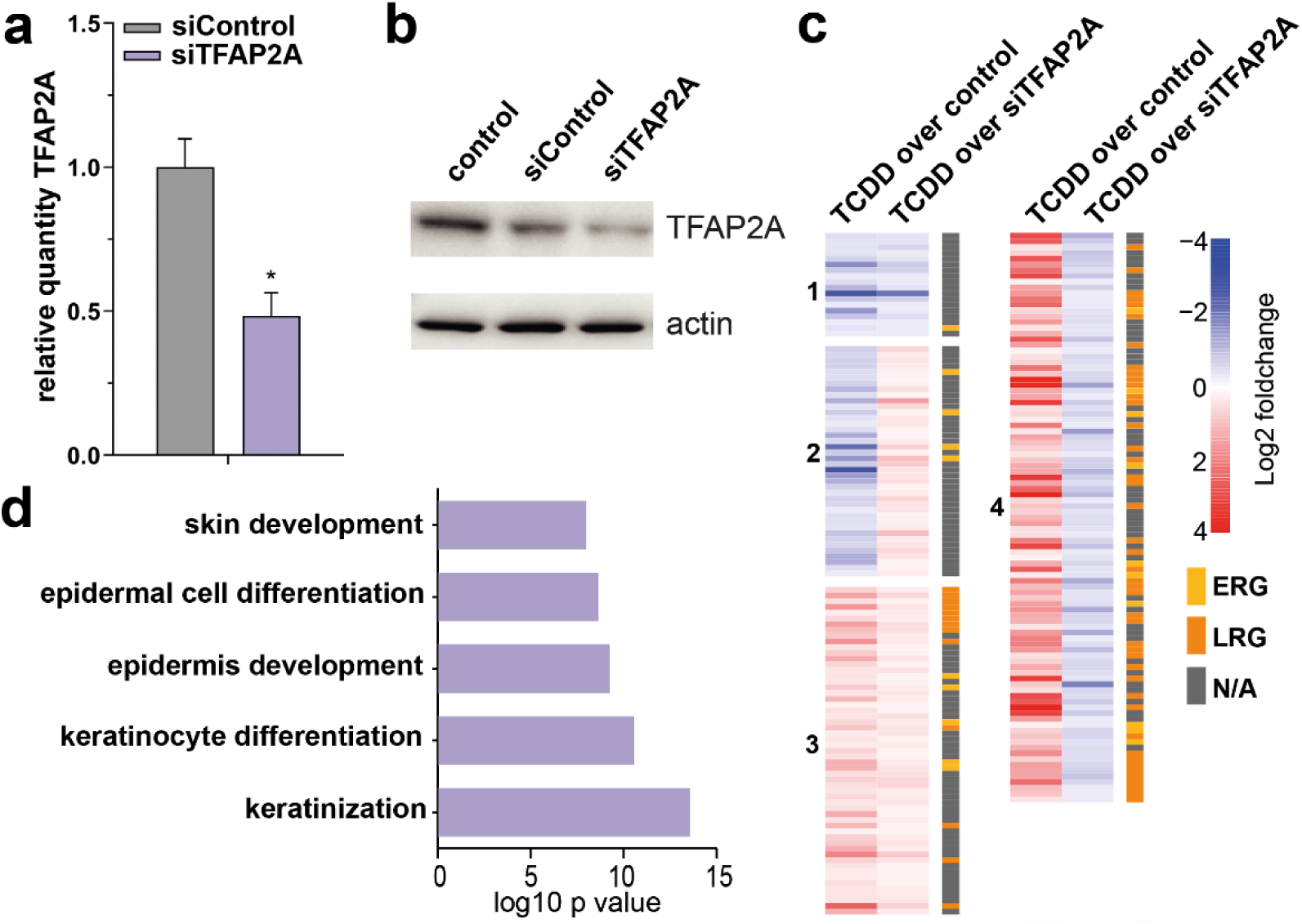
AHR-TFAP2A axis controls the epidermal differentiation program a. Validation of *TFAP2A* knockdown by RT-qPCR, normalized to reference gene *hARP*. Data shown as mean ±SEM, N=3 technical replicates, unpaired T-test. **b** Western blot validation of TFAP2A knockdown. Actin was used as a loading control for the quantification of TFAP2A protein levels. **c** Heatmap of differentially expressed genes (p value <0.05), accompanied by ERG (yellow) or LRG (orange) nomination. Z-score was calculated based on log10 (FPKM+0.01) of each gene. **d** GO annotation of genes in cluster 4 of the heatmap.

To investigate whether TFAP2A directly regulates these genes, we sought for the TFAP2A binding motif near promoter and enhancers of genes in cluster 4. We found that 64 out of 99 genes have a TFAP2A binding motif at their promoter regions while all 99 genes have TFAP2A motif at their enhancer regions (Supplementary Data 5). These results indicate that TFAP2A likely regulate these cluster 4 genes directly, and support the notion that AHR controls keratinocyte differentiation through activation of TFAP2A.

Finally, the importance of the identified AHR-TFAP2A axis in keratinocyte differentiation was investigated by knocking out TFAP2A using CRISPR/Cas9 on immortalized N/TERT-2G keratinocytes. Clonal homozygous TFAP2A knockout (ΔTFAP2A) N/TERT-2G keratinocytes were generated, grown in monolayer cultures, and treated with TCDD for up to 72 h, similar to the conditions of the previously described AHR-activated siRNA experiment. The upregulation of the AHR target gene *CYP1A1* by TCDD was not altered in ΔTFAP2A keratinocytes, indicating that *CYP1A1* is not regulated through TFAP2A (Fig. 5a). However, differentiation-related AHR-responsive genes, e.g., *IVL*, *SPRR1A/B*, *SPRR2*, and *MMP1,* of which expression could be induced by TCDD in wildtype keratinocytes, were not upregulated in ΔTFAP2A keratinocytes (Fig. 5b). The expression patterns of these genes upon TCDD treatment with TFAP2A depletion are consistent with those observed from the siTFAP2A experiment (cluster 4)(Fig. 5b, compared to untreated condition, only 72 h timepoint shown). Consistently, the lack of mRNA upregulation induced by TCDD in ΔTFAP2A keratinocytes was observed for many other epidermal differentiation genes detected by RT-qPCR, e.g., *PRR9*, *DSG1*, *DSC1*, *S100A8*, *TRPV3*, and *TGM3* (Suppl. Fig. 1a). Importantly, already at baseline, ΔTFAP2A keratinocytes showed significantly less expression of these genes as compared to wildtype N/TERT-2G keratinocytes (Fig. 5c), indicating that loss of TFAP2A is not adequately compensated. Interesting to note, expression of *AHR* was not hampered in ΔTFAP2A keratinocytes (Suppl. Fig. 1b), implying that TFAP2A is not part of a self-regulating AHR signaling feedback loop. In summary, these data demonstrate that TFAP2A is an indispensable regulator in the molecular cascade of AHR-mediated keratinocyte differentiation. although it is unlikely that TFAP2A is involved in other AHR-mediated biological processes, such as xenobiotic metabolism where *CYP1A1* is a target.

**Fig. 5.**
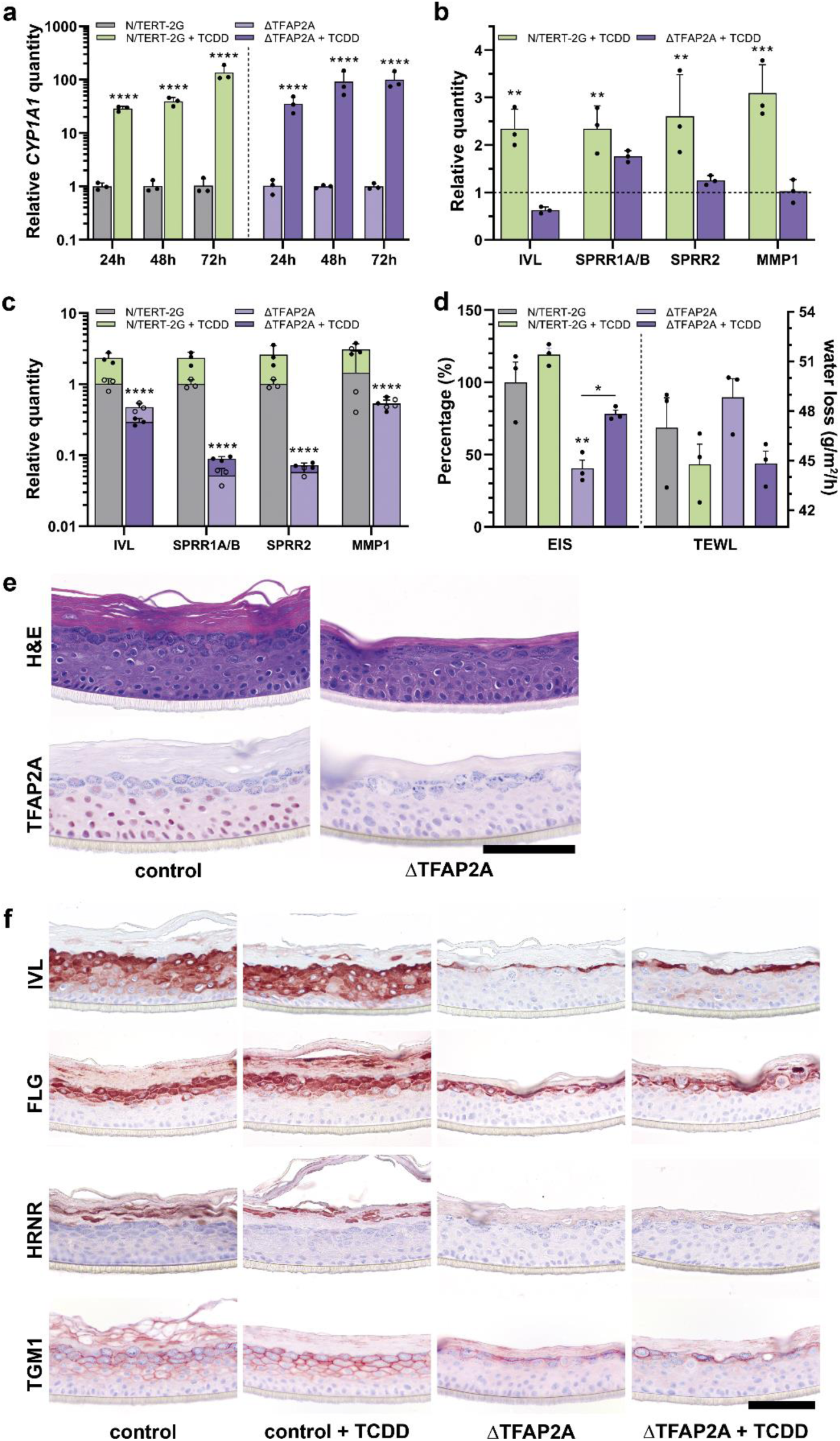
AHR-TFAP2A axis in keratinocyte differentiation and function. a. Monolayer N/TERT-2G and ΔTFAP2A were treated with TCDD for up to 72 h and AHR activation was validated by *CYP1A1* RT-qPCR. Data are shown as mean ±SEM, N=3 technical replicates, two-way ANOVA. **b** RT-qPCR analysis of several genes from cluster 4 (Fig. 4c) displays AHR-dependent induction in the N/TERT-2G keratinocytes but not in ΔTFAP2A keratinocytes. Data are compared to their respective untreated condition and shown as mean ±SEM, N=3 technical replicates, two-way ANOVA. **c** In addition, RT-qPCR analysis shows significant reduction of basal gene expression in ΔTFAP2A keratinocytes regardless of AHR activation. TCDD treatment data (closed circles) shown superimposed on untreated data (open circles). Data depicted as mean ±SEM, N=3 technical replicates. **d** Functional skin barrier analyses EIS and TEWL on HEEs and ΔTFAP2A-HEEs displays reduced electrical impedance and increased transepidermal water loss, indicating a reduced barrier functionality. Barrier functionality is improved by TCDD treatment, as EIS increases and TEWL reduces. Data shown as mean ±SEM, N=3 technical replicates, one-way ANOVA. TEWL differences are not significant due to variation in the untreated HEEs. Full EIS spectrum in Supplementary Fig. 1c. **e** Immunohistochemistry confirms the complete loss of TFAP2A expression and **f** indicates the reduction of IVL, FLG, HRNR, and TGM1 expression, while TCDD treatment marginally upregulates the expression of IVL and FLG. Scale bar = 100 µm.

ΔTFAP2A organotypic human epidermal equivalents (ΔTFAP2A-HEEs) were generated to examine whether TFAP2A knockout and accompanied loss of keratinocyte differentiation gene expression gives rise to morphological changes and epidermal barrier defects. Quantitative epidermal barrier properties were analyzed by electrical impedance spectroscopy (EIS) and transepidermal water loss (TEWL) (Fig. 5d, complete EIS spectrum in Suppl. Fig. 1c). ΔTFAP2A-HEEs showed reduced electrical impedance, indicating functional skin barrier defects of ΔTFAP2A-HEEs, which agrees with the altered keratinocyte differentiation gene expression. Of note, we observed statistically significant improvement in the EIS values upon AHR activation by TCDD in ΔTFAP2A-HEEs, which was corroborated by a non-significant trend of reduction in TEWL. The loss of TFAP2A expression was confirmed by immunochemistry staining (Fig. 5e), which coincided with altered epidermal morphology, e.g., less flattened keratinocytes, less *stratum granulosum*, and thinner *stratum corneum*, likely caused by aberrant differentiation. Indeed, downregulated protein expression of a panel of important terminal differentiation proteins was detected, including IVL, FLG, HRNR, and TGM1 was found in ΔTFAP2A-HEEs (Fig. 5f).

These data suggest that loss of TFAP2A can partially be alleviated upon AHR activation, presumably by other AHR-induced ERGs (e.g., *OVOL-1*^41^, fold change 2.29 (Supplementary Data 1)) that cooperate in the terminal differentiation program.

## Discussion

In this study, we aimed to elucidate the signaling cascades by which the AHR exerts transcriptional regulation of terminal differentiation and skin barrier formation. We combined transcriptomic and epigenomic analyses to characterize the temporal gene regulatory events following AHR activation using keratinocytes as a model system for barrier epithelia. We identified that in a temporal distinct early response, AHR directly regulates the expression of several TFs known to be important for skin development and keratinocyte differentiation, e.g., *TFAP2A*^42–44^. Furthermore, we found that TFAP2A directly enhances epidermal differentiation as a secondary response to AHR activation and thereby contributes to skin barrier integrity. Low-level activation of AHR by endogenous, circulating, weak AHR agonists might drive the TFAP2A-mediated keratinocyte differentiation *in vivo*. As such, these findings further elucidate the molecular mechanisms of action by which AHR induces its target effects.

Among the many biological roles that AHR has been associated with in the skin, our study specifically unravels the molecular mechanism behind AHR-mediated keratinocyte differentiation. We identified distinct early and late responses upon AHR activation where TFs activated during the early response such as TFAP2A regulate keratinocyte differentiation genes in the late response. In addition, we demonstrate that AHR activation leads to enhancer dynamics that distinguish direct targets from secondary effects. The AHR:ARNT binding motif was significantly enriched in dynamic enhancers, which were pre-established open chromatin regions with visible H3K27ac signals already before the treatment started. Thus, instead of establishing *de novo* enhancers, like pioneer transcription factors (e.g., p63) that orchestrate the cell-type-specific enhancer landscape^45,46^, AHR seems to exploit a pre-specified landscape of targets. This enables a swift response towards external threats through regulation of canonical pathways like the cytochrome P450 pathway and mitogen-activated protein kinase (MAPK)^47^. Enhancers that showed dynamic H3K27ac signals at later time points were located near genes involved in ‘keratinocyte differentiation’, of which activation represents secondary effects of AHR activation. Interestingly, many of these genes are considered AMPs, consistent with our and others’ recent findings that AHR activation in keratinocytes induces AMP expression^16,48^. It is important to note that AHR direct targets that have AHR:ARNT motif-containing enhancers nearby, e.g., *CYP1A1*, are not all regulated by TFs such as TFAP2A, indicating the specificity of AHR action in different biological processes. In addition, immune system related functions appear to be associated with both pre-established and dynamic chromatin regions, suggesting that different immune genes are either induced or repressed at different time points upon AHR activation (Supplementary Data 2). This intriguing complexity and in the temporal cooperation between different immune pathways in response to environmental threats are subject to future research and may shed further light on the Janus-faced role of AHR^49^.

Unlike AHR binding profiles in cancer cell lines that contain thousands of AHR binding sites (including TFAP2A)^50^, our AHR ChIP-seq in keratinocytes yielded ‘only’ 57 AHR binding site, probably due to the transient binding nature of AHR upon ligand activation in normal cells. The validation of several sites by ChIP-RT-qPCR strengthens our confidence that these are genuine AHR-bound regions having biological relevance. The use of cancer cell lines (e.g., HaCaT keratinocytes) in this field of research may thus overestimate the number of target genes that are actually bound by AHR under physiological conditions.

The similarity observed between TCDD and CT treatment *in vitro* is striking, when considering that these are on the opposite sides of the health spectrum: TCDD being highly toxic while CT is used as a dermatological therapy for psoriasis and atopic dermatitis^14,16,51-53^. Both ligands activated AHR similarly, provoking an adjective transcriptional response in keratinocytes. However, it is important to realize that the short-term effects of AHR activation in an experimental *in vitro* system do not take into account the ligand metabolism, degradation and elimination that would normally occur *in vivo*. TCDD’s extreme long half-life (not being a substrate for xenobiotic metabolism) and systemic exposure has devastating chronic effects through sustained AHR activation^54^. In contrast, CT is a highly complex mixture of many different chemicals that could counter-act or compensate for the agonistic effects and is given only localized and periodically to patients. This raises an interesting question on the proposed AHR ligand promiscuity at the molecular level^55^. The dosage and half-life of AHR ligands and thus strength and duration of AHR activation may determine the biological effect. Whether AHR ligands are stable, rapidly metabolized, or whether secondary metabolites are involved in activities independent of AHR signaling pathways requires further investigation^56^. Herein, timing seems to be of utmost importance and time-course global gene expression profiling in vivo is an essential next step to evaluate AHR activation and to dissect this regulatory cascade to greater detail.

Over the years, evidence has grown that serum levels of dietary-derived or microbiota-derived components can activate the AHR in several barrier organs *in vivo*. For example, indole-3-carbinole (I3C) can robustly activate the AHR in the intestine^57^ whereas tryptophan metabolites can regulate AHR activation in the skin (e.g., FICZ^58^, kynurenine^59^, and kynurenic acid^60^). This implies that dietary intervention can be helpful in controlling AHR activation and thus support TFAP2A-mediated skin barrier integrity.

To conclude, our findings indicate that activation of AHR triggers a regulatory cascade mediating keratinocyte differentiation and this cascade relies on TFs such as TFAP2A that play an intermediate but indispensable role. Our discovery on the AHR-TFAP2A axis exemplifies how environmental factors can dictate the terminal differentiation process, and unveil alternative routes and targets that may be hijacked to foster barrier formation and repair in the skin (and presumably other barrier organs) without the need for AHR activation per se.

## Acknowledgements

This research has been a long endeavor with many collaborative efforts on the various aspects of the experimental work and data analysis. We are grateful for all funders throughout the years: Radboud Institute for Molecular Life Science (RIMLS; JSc and EB), National Institutes of Health (ES028244); Dutch Research Council VENI-grant 91616054 (EB), Chinese Scholarship Council grant 201406330059 (JQ), and LEO Foundation grant LF18068 (PZ and EB). We thank all members from the Departments of Dermatology, Molecular Developmental Biology, and Molecular Biology for discussion and suggestions on the project. We especially thank Chet Loh for discussion and useful input. We thank Eva Janssen-Megens, Siebe van Genesen and Rita Bylsma for operating the Illumina analyzer. We thank the ENCODE Consortium for sharing their data and Gary H. Perdew for the critical reading of our manuscript.

## Author contributions

JSm, JQ, HZ, and EB conceived and designed the study and experiments and supervised the data analysis. JSm, JQ, NB, DRO, and IVW performed the wet-lab experiments. The AHR knockout line was generated by NB, the TFAP2A knockout line by JSm. JSm, JQ, FP, NB, SH, HZ, and EB analyzed the data. For the omics-analysis in particular, FP performed the siRNA transcriptome data analysis, JQ, SH and HZ were responsible for the data integration. JSm, JQ, FP, NB, JSc, PZ, HZ, and EB wrote and/or revised the manuscript.

## Material and methods

### Cell culture and drug treatment

Human abdominal or breast skin was obtained from plastic surgery procedures after informed consent and in line with the principles and guidelines of the Declaration of Helsinki. Skin biopsies were taken and human primary keratinocytes were isolated as previously described ^61^ and stored in liquid nitrogen until further use. Human primary keratinocytes were cultured in Keratinocyte Basal Medium (KBM, Lonza #CC-4131) supplemented with 0.4% (vol/vol) bovine pituitary extract, 0.5 μg/mL hydrocortisone, 5 μg/mL insulin and 10 ng/mL epidermal growth factor (Lonza #CC-4131). Medium was refreshed every other day until near confluency before treatment commencement. Dimethylsulfoxide (DMSO) was purchased from Merck (Darmstadt, Germany), 2,3,7,8-Tetrachlorodibenzo-p-dioxin (TCDD) was purchased from Accustandard (New Haven, CT, USA), and coal tar (CT) was purchased from Fagron BV (Capelle aan den IJssel, The Netherlands). Cells were treated with either DMSO (0.1% vol/vol), CT (4 µg/mL), or TCDD (10 nM). Total RNA was collected for RNA-seq and qPCR-based validation purposes. Chromatin was harvested for ChIP-seq experiments. Lysates containing proteins were harvested for western blotting purposes. No mycoplasma contaminations were found during cell culture.

### N/TERT-2G culture and human epidermal equivalent (HEE) generation

Human N/TERT keratinocyte cell line N/TERT-2G, purchased from J. Rheinwald laboratory (Harvard Medical School, Boston, MA, USA), was cultured in Epilife medium (MEPI500CA, ThermoFisher Scientific, Waltham, MA, USA), complemented with human keratinocyte growth supplement (S0015, ThermoFisher Scientific) and 1% penicillin/streptomycin (P4333, Sigma-Aldrich, Saint-Louis, MO, USA). Human epidermal equivalents (HEEs) were generated as previously described^62^, with minor adjustments. Briefly, inert Nunc cell culture inserts (141002, ThermoFisher Scientific) were coated with rat tail collagen (100 µg/mL, BD Biosciences, Bedford, USA) at 4°C for 1 hour. 1.5×10^5^ N/TERT-2G keratinocytes (either wildtype, ΔAHR, or ΔTFAP2A keratinocytes) were seeded on the transwells in 150 µL Epilife medium (ThermoFisher Scientific) supplemented with 1% penicillin/streptomycin (Sigma-Aldrich) in a 24 wells format. After 48 h, cultures were switched to a mixture of CnT-PR-3D medium (CELLnTEC, Bern, Switzerland) and DMEM medium (60:40 (v/v)) without penicillin/streptomycin for 24 h and then cultured at the air-liquid interface for an additional ten days. Culture medium was refreshed every other day until harvesting at day ten of the air-exposed phase.

### Western blotting and Immunofluorescence

Cell lysates of human primary keratinocytes were collected after treatment using RIPA lysis buffer. Afterwards, the lysates were sonicated (10x 5s on/off) and the samples were loaded onto SDS PAGE gel and transferred to PVDF membranes using the NuPAGE system (Life technologies) and visualized using SuperSignal West Femto Maximum Sensitivity Substrate (ThermoFisher, #34095). For analysis of AHR translocation to the nucleus, direct immunofluorescence (IF) labeling was performed as described^14^. Antibodies for western blotting and IF are listed in Table 5.

### Transcriptional analysis by quantitative real-time PCR

Total RNA was isolated using the Favorprep total tissue RNA kit (Favorgen Biotech, Taiwan), according to the manufacturer’s protocol. cDNA was generated after DNase treatment and used for quantitative real-time PCR (RT-qPCR) by use of the MyiQ Single-Colour Real-Time Detection System (Bio-Rad laboratories, Hercules, CA, USA) for quantification with Sybr Green and melting curve analysis. Primers (Table 2) were obtained from Biolegio (Nijmegen, The Netherlands) or Merck. Target gene expression levels were normalized to the expression of human acidic ribosomal phosphoprotein P0 (RPLP0). The relative expression levels of all genes of interest were measured using the 2-ΔΔCT method^63^.

**Table 2:**
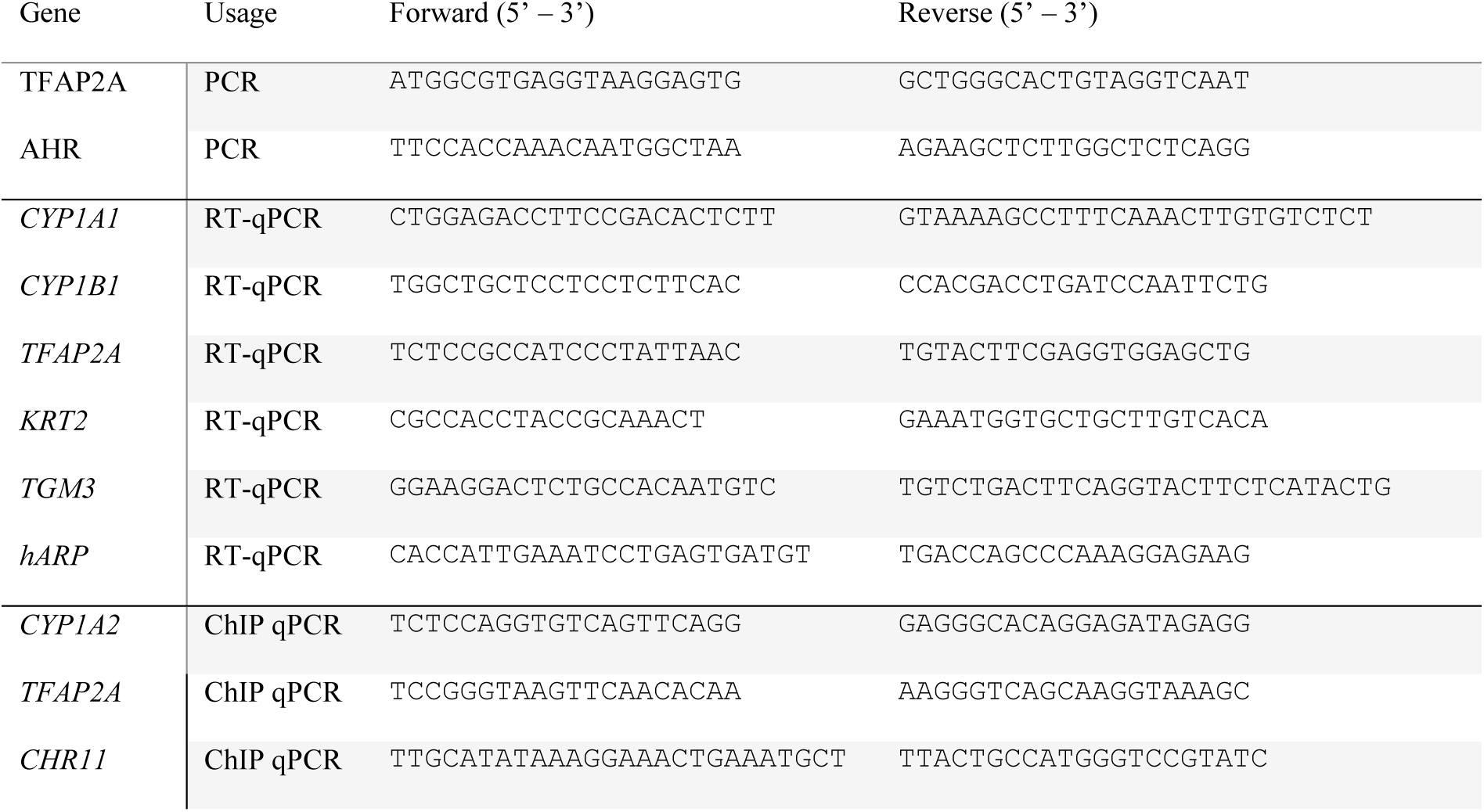
PCR, RT-qPCR and ChIP qPCR primers

### RNA sequencing and analysis pipeline

RNA sequencing was performed as described previously^45^ with the starting material of 500 ng total RNA, to obtain double-strand cDNA (ds-cDNA). After purification with the MinElute Reaction Cleanup Kit (Qiagen #28206), 3 ng ds-cDNA was processed for library construction using KAPA Hyper Prep Kit (Kapa Biosystems #KK8504) according to the standard protocol except that a 15-min USER enzyme (BioLab # M5505L) incubation step was added before library amplification. The prepared libraries were quantified with the KAPA Library Quantification Kit (Kapa Biosystems #KK4844), and then sequenced in a paired-ended manner using the NextSeq 500 (Illumina) according to standard Illumina protocols.

Sequencing reads were aligned to human genome assembly hg19 (NCBI version 37) using STAR 2.5.0^64^ with default options. Briefly, STAR has the option to generate in-house reference genome from the genome fastq file. In this study, hg19 genome was used to generate the in-house reference genome with the following command: STAR --runThreadN 8 --runMode genomeGenerate --genomeDir directory/ --genomeFastaFiles hg19.fa --sjdbGTFfile Homo_sapiens.GRCh37.75.gtf --sjdbOverhang 100. Then STAR was run and it automatically generated read-counts directly. For data visualization, wigToBigWig from the UCSC genome browser tools was used to generate bigwig files and uploaded to UCSC genome browser. Genes with the mean of DESeq2-normalized counts (“baseMean”)> 10 were considered to be expressed. Differential gene expression (adjusted P value <0.05) and principal-component analysis were performed with the R package DESeq2 using read counts per gene^65^. Hierarchical clustering was performed based on log10 (FPKM+0.01). Functional annotation of genes was performed with DAVID^66^. For the experiments containing siRNAs, read counts were generated as described above. Differential expression analysis was performed using R community-created R packages stringr^67^ and dplyr^68^, and the DESeq2 package with normalization on siRNA treatment (DESeq design = siRNA). Read counts from control and TCDD treated samples at the 24 h stimulation time point were re-analyzed in a separate DESeq2 differential expression analysis (DESeq design = treatment). Significant differentially expressed genes overlapping between both experiments (Benjamini & Hochberg adjusted p-value < 0.05)^69^ were visualized in a heatmap using the ComplexHeatmap package^70^. Gene Ontology analysis of interesting groups was performed using clusterProfiler^71^. Identification of TFs was performed as described before^72^.

### ChIP sequencing and analysis pipeline

Chromatin for ChIP was prepared as previously described^73,74^. ChIP assays were performed following a standard protocol^75^ with minor modifications. On average, 0.5M keratinocytes were used in each ChIP. For histone mark H3K27ac, 2x ChIP reactions were pooled to prepare 1x ChIP-seq sample; for AHR, 4x ChIP reactions are pooled to prepare 1 ChIP-seq sample. Antibodies against H3K27ac (Diagenode C15410174) and AHR (Santa Cruz Biotechnology Inc. Santa Cruz, CA, USA, sc-5579) were used in each ChIP assay. Resulted DNA fragments from four independent ChIP assays were purified and subjected to a ChIP-qPCR quality check. Afterwards 5ng DNA fragments were pooled and proceeded on with library construction using KAPA Hyper Prep Kit (Kapa Biosystems #KK8504) according to the standard protocol. The prepared libraries were then sequenced using the NextSeq 500 (Illumina) according to standard Illumina protocols.

Sequencing reads were aligned to human genome assembly hg19 (NCBI version 37) using BWA^76^. Mapped reads were filtered for quality, and duplicates were removed for further analysis. In addition. The bamCoverage script was used to generate and normalize bigwig files with the RPKM formula. The peak calling was performed with the MACS2^77^ against a reference input sample from the same cell line with standard settings and a q value of 0.05. Only peaks with a p value < 10e-5 were used for differential analysis with MAnorm^78^. Association of peaks to genes and associated GO annotation were performed with GREAT^79^, with the ’single nearest gene within 1 Mb’ association rule. *P* values were computed with a hypergeometric distribution with FDR correction. k-means clustering and heat map and band plot generation were carried out with a Python package fluff^80^. HOMER (http://homer.salk.edu/homer/motif/) was used for motif scan against corresponding background sequences. One thing needs to be mentioned is that we overlapped dynamic enhancers with published DNAse I hypersensitivity sites to narrow down regions for motif scan.

### ATAC-seq and motif analysis

ATAC-seq dataset (GSE123711) was downloaded and used for motif enrichment analysis as described before^81^. Briefly, ATAC-seq peaks within TSS-1Kb to TSS+0.5Kb were defined as promoter regions, whereas ATAC-seq peaks TSS-1Mb to TSS+1Mb were defined as enhancer regions. Differential motif analysis and TFAP2A motif scan within promoter regions and enhancer regions were separately performed using HOMER tool using default parameters (http://homer.salk.edu/homer/motif/).

### siRNA knockdown

Human primary keratinocytes were grown to 10-15% confluency before 500 nM of Accell human SMARTpool gene targeting or non-targeting siRNA (Dharmacon, Lafayette, CO, USA) was added for 48 h. Culture medium was subsequently refreshed and supplemented with siRNA for another 48 h. Keratinocyte were thereafter allowed to differentiate for 24 h in the presence of TCDD and were harvested for transcriptional analysis and western blotting as described above. siRNA SMARTpools include: Accell Human TFAP2A (7020) SMARTpool (#E-006348-00) and Accell Non-targeting Control Pool (#D-001910-10).

### Single guide RNA design, single strand donor oligonucleotide and synthetic Cas9

Synthetic sgRNAs to knockout *AHR* and *TFAP2A* gene, and purified Edit-R Cas9 nuclease protein (NLS, #CAS11200) were bought from Invitrogen (Waltham, MA, USA) and IDT Technologies (Coralville, IA, USA), respectively. See Table 3 for details on the sgRNAs used.

**Table 3:**
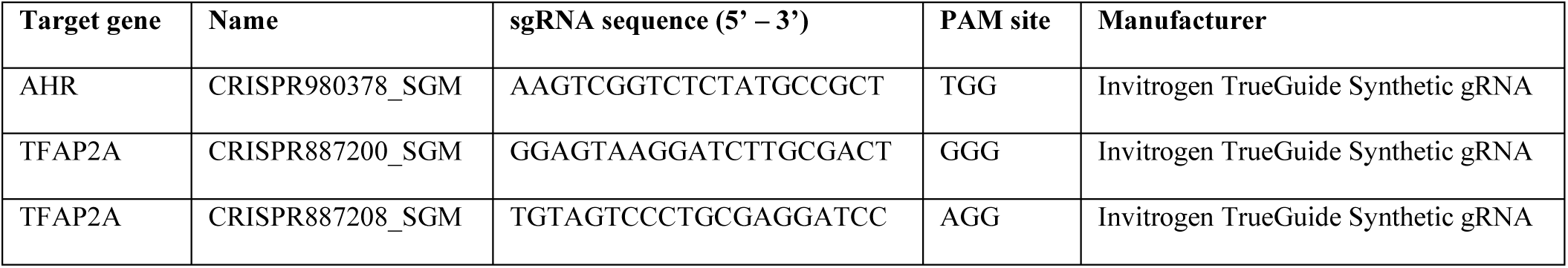
Sequences of the sgRNAs.

### Electroporation of ribonucleoprotein (RNP) complexes and analysis of editing efficiency

N/TERT-2G keratinocytes were electroporated using the NEON transfection system 10µL kit (ThermoFisher Scientific). N/TERT-2G keratinocytes were detached from culture plastic and washed twice with dPBS (without Ca^2+^ and Mg^2+^) as described above. Meanwhile, per electroporation condition, synthetic sgRNA (300ng) and Cas9 (1.5 µg) were incubated with 5 µL resuspension buffer R for 20 min before adding 1×10^5^ N/TERT-2G keratinocytes. After mixing the cell suspension, the cells were electroporated using 1 pulse of 1700V for a duration of 20ms before immediate seeding in a pre-warmed 6 well plate. DNA was isolated using the QIAamp DNA blood mini kit (51106, Qiagen, Hilden, Germany) according to manufacturer’s protocol after a couple of days and CRISPR/Cas9 induced editing efficiency was analyzed by PCR and separation of amplicon on 2% agarose gel containing 1:10.000 GelRed nucleic acid gel stain (41003, Biotium Inc., Fremont, CA, USA). Amplicons were purified by MinElute Gel extraction kit (28606, Qiagen) using the manufacturers protocol and sanger sequenced to assess editing efficiency. Sanger sequencing reads were analyzed using the Inference of CRISPR edits (ICE) webtool (ice.synthego.com, v3, Synthego Corporation, Menlo Park, CA, USA). See Table 4 for details on the PCR primers used.

**Table 4:**
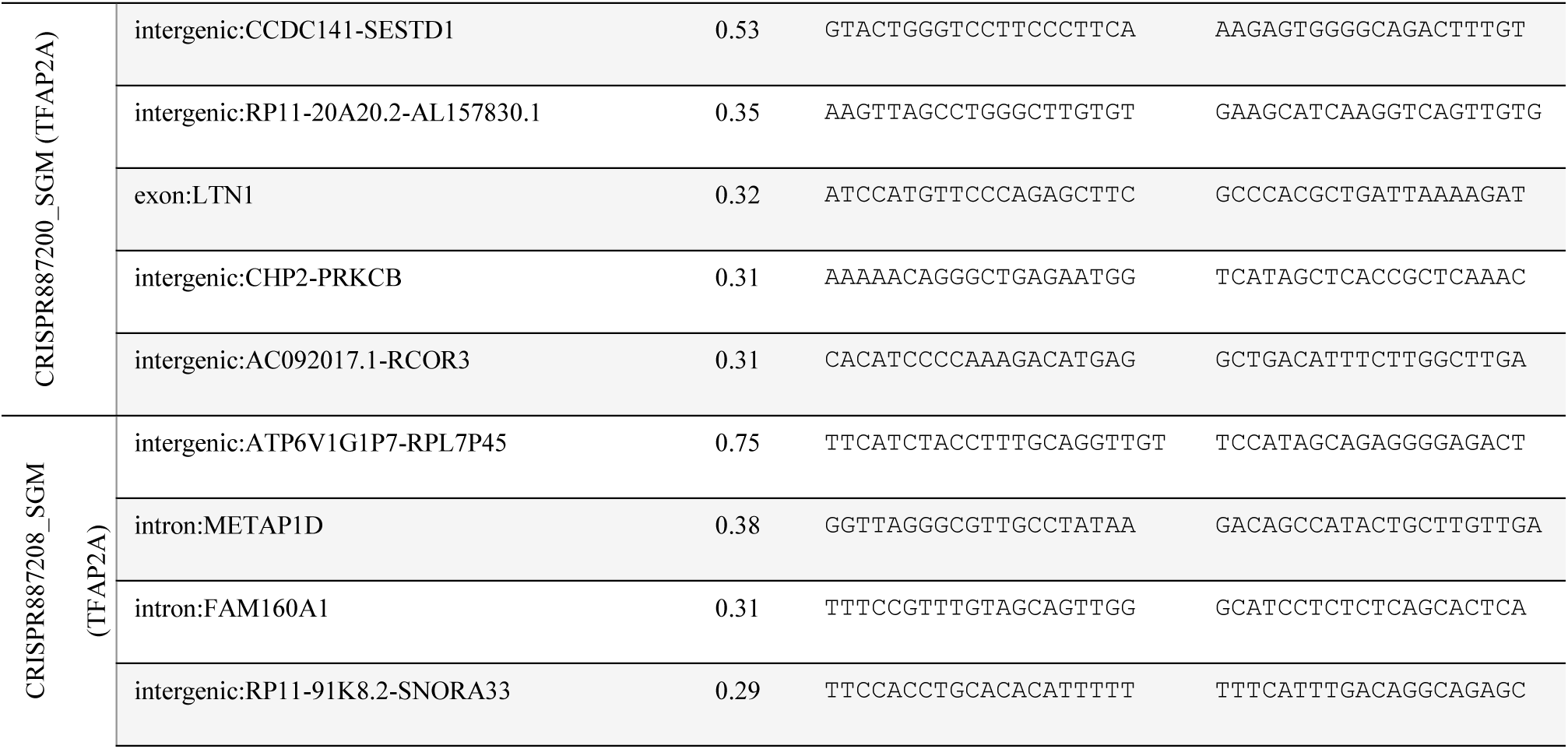
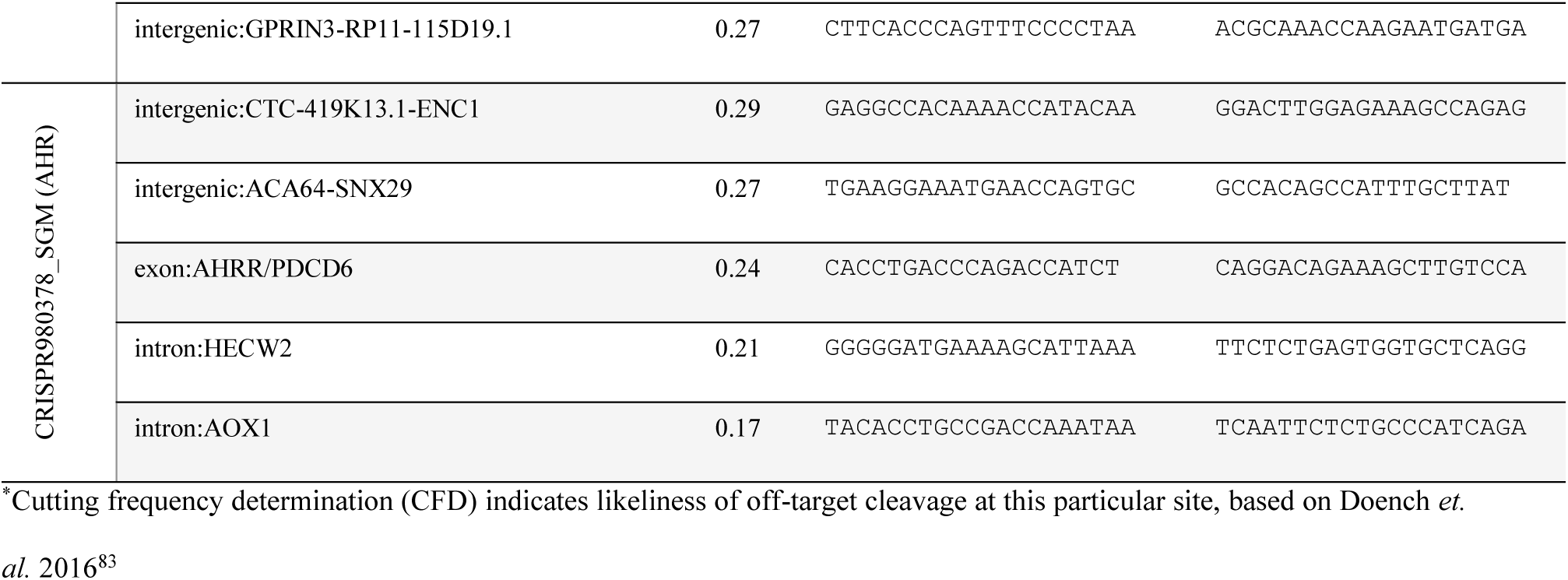
Regular PCR primers for predicted off-target site analysis

### Generation of clonal ΔAHR and ΔTFAP2A N/TERT keratinocytes

N/TERT-2G keratinocyte cell pools carrying AHR or TFAP2A knockouts were diluted to seed one cell per well (∼600 cells per 60 mL of Epilife medium, 100µL per well) into 6×96 well plates and allowed to grow for one week before refreshing the medium. After another week of culture, cells were passaged, as described above, into 24 well plates, 6 wells plates, T25 flasks, and T75 flasks subsequently before freezing them. Cell clonality was assessed by sanger sequencing and analyzing genomic DNA at the targeted locus with help of the ICE webtool (ice.synthego.com, v3, Synthego Corporation).

### *In silico* search for potential off-target effects

CRISPOR (version 4.99)^82^ was used to search for potential off-target effects dependent on the *streptococcus pyogenes* derived Cas9 (SpCas9) PAM site (5’-NGG-3’), target genome (homo sapiens GRCh38/hg38) and our specific guide RNA selection. Using genomic DNA of the N/TERT-2G keratinocyte knockout clones, the top-5 off-target sites for all guide RNAs were amplified by PCR and analyzed by sanger sequencing to assure no off-target mutations occurred. See Supplementary Data 6 for off-target analysis results and Table 4 for details on the PCR primers used for off-target analysis.

### Epidermal barrier measurements TEWL and EIS

Epidermal barrier capabilities of epidermal equivalent cultures were studied by use of transepidermal water loss (TEWL) measurements and electrical impedance spectroscopy (EIS). After habituation of the cultures to room temperature, TEWL was measured using the Aquaflux AF200 (Biox, UK) on day 10 of the air-exposed phase of the HEE culture. TEWL was measured in triplicate in wildtype N/TERT-2G keratinocyte and ΔTFAP2A keratinocyte HEEs. Significance was assessed using one-way ANOVA with multiple comparisons correction (Tukey). EIS was measured using the real-time impedance detector Locsense Artemis (Locsense, Enschede, The Netherlands) with the SmartSense lid for monitoring cells in conventional transwell plates with inserts. Impedance (Ω) measurements were performed on day 10 of the air-exposed phase of the HEE culture after habituation of the HEE cultures to room temperature. EIS was measured in triplo on wildtype N/TERT-2G keratinocytes and ΔTFAP2A keratinocyte HEEs. After calibration, continuous impedance (Ω) was measured using standard settings e.g., sweeping frequency from 10Hz to 100.000Hz. Afterwards, measured impedance was corrected with blank impedance measurements per electrode and corrected for the size of the culture insert (0,47cm^2^), resulting in impedance per cm^2^ values (Ω/cm^2^). Significance was assessed using one-way ANOVA with multiple comparisons correction (Tukey).

### Morphological and immunohistochemical analysis of HEEs

HEEs were fixed in 4% formalin solution for 4 h and subsequently embedded in paraffin. 6 µm sections were stained with hematoxylin and eosin (H&E, Sigma-Aldrich) or processed for immunohistochemical analysis. Sections were blocked for 15 min with 5% normal goat or horse serum in phosphate-buffered saline (PBS) and subsequently incubated with the specific antibodies for 1 h at room temperature. Next, a 30 min incubation step with biotinylated horse anti-mouse, or goat anti-rabbit (Vector Laboratories, Burlingame, USA) was performed, followed by a 30 min incubation with avidin-biotin complex (Vector Laboratories). The peroxidase activity of 3-Amino-9-ethylcarbazole (AEC) was used to visualize the protein expression and the sections were mounted using glycerol gelatin (Sigma-Aldrich). See Table 5 for details on the antibodies used for immunofluorescence, western blot, and immunohistochemistry.

**Table 5:** Antibodies used in immunohistochemistry and western blot

| Purpose* | Antibody | Manufacturer and catalog number | Dilution |
| --- | --- | --- | --- |
| IF | Rabbit anti-AHR | Santa Cruz Biotechnology, SC-5579 | 1:200 |
| IHC | Mouse anti-CYP1A1 | Santa Cruz Biotechnology, SC-25304 | 1:25 |
| WB / IHC | Mouse anti-TFAP2A | Invitrogen, MA1-872 | WB 1:300<br>IHC 1:50 |
| WB | Mouse anti-β-Actin, AC-15 | Merck, Darmstadt, Germany | 1:100.000 |
| IHC | Mouse anti-FLG | Leica Biosystems, Newcastle, UK | 1:100 |
| IHC | Rabbit anti-HRNR | Sigma-Aldrich, HPA031469 | 1:500 |
| IHC | Mouse anti-IVL, Mon150 | Van Duijnhoven *et al*<sup>84</sup> | 1:20 |
| IHC | Rabbit anti-Ki67 | Abcam, Cambridge, UK, ab16667 | 1:50 |
| IHC | Horse anti-mouse, biotinylated | Vector Laboratories, BA-200-1.5 | 1:200 |
| IHC | Goat anti-rabbit, biotinylated | Vector Laboratories, BA-5000-1.5 | 1:200 |

\*IF = immunofluorescence; WB = western blot; IHC = immunohistochemistry

### Statistics and reproducibility

Data set statistics were analyzed using the GraphPad Prism software. Differences under p value < 0.05 were considered statistically significant, ns p value > 0.05, * p value <0.05, ** p value <0.01, *** p value <0.001, **** p value <0.0001. Gene expression analysis by RT-qPCR was performed in biological duplicates (at least n=3); data are shown as mean ± standard error of the mean unless otherwise specified. Statistics was performed on dCT values using one-way ANOVA with multiple comparison correction (Tukey). Other statistical methods used are specified in the methods sections.

## Data availability

All raw sequencing files including RNA-seq and ChIP-seq analyses generated in this study have been deposited in the GEO database with the accession number GSE226047.

---

**Supplementary Fig. 1.**
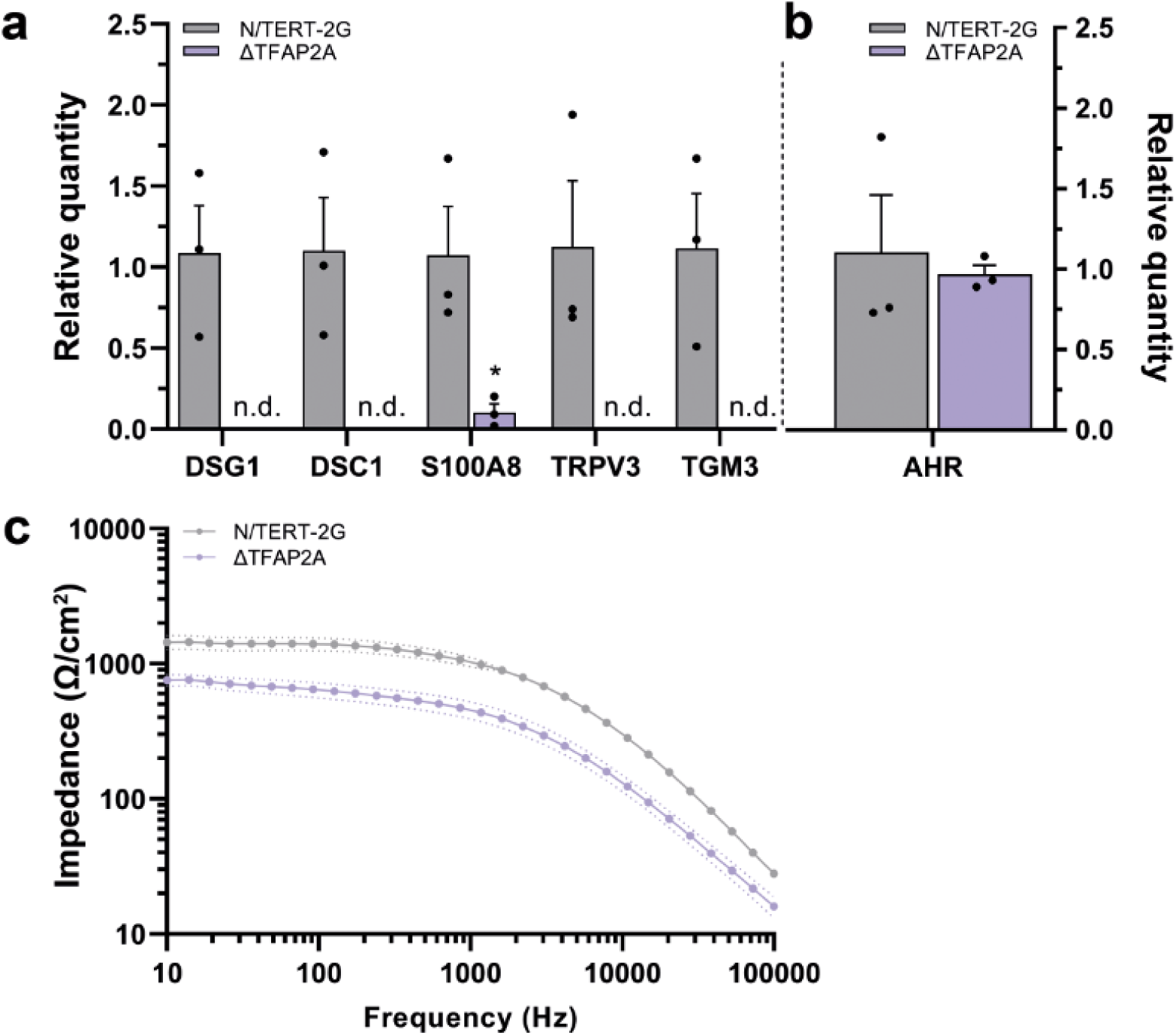
**a** RT-qPCR analysis of monolayer untreated N/TERT-2G and ΔTFAP2A keratinocytes indicates that loss of TFAP2A results in severe downregulation of several epidermal differentiation genes (n.d. = nondetectable). Data are compared to N/TERT-2G keratinocytes and shown as mean ±SEM, N=3 technical replicates, one-way ANOVA. **b** *AHR* expression is not changed in monolayer ΔTFAP2A keratinocytes as shown per RT-qPCR analysis. Data shown as mean ±SEM, N=3 technical replicates, one-way ANOVA. **c** Full EIS spectrum of HEEs and ΔTFAP2A-HEEs (from Fig. 5f) showing reduced electrical impedance of ΔTFAP2A-HEEs. Data shown as mean ±SEM, N=3 technical replicates.

